# Expression Pattern of Epilepsy Associated Genes as a Common Pathological Signature in PTE and SE models and possible therapeutics via an in-silico approach

**DOI:** 10.64898/2026.04.23.720518

**Authors:** J. Tyagi, S. Kumari, C. Prakash, S. Saran, B. Kumar Biswal, D. Sharma

**Affiliations:** Neurobiology Laboratory, School of Life Sciences, Jawaharlal Nehru University, New Delhi-110067, India; Structural and Functional Biology Laboratory, National Institute of Immunology, New Delhi-110067, India; Cell and Developmental Biology Laboratory, School of Life Sciences, Jawaharlal Nehru University, New Delhi-110067, India

**Keywords:** Seizures, post-traumatic epilepsy, status epilepticus, electrophysiology, natural bioactive compounds

## Abstract

The transition from an initial brain insult to chronic epilepsy remains a critical challenge in clinical neuroscience, often involving complex molecular shifts that are difficult to capture through single-gene analysis. This study investigated the existence of a convergent pathway of genomic instability across two distinct etiological models: iron-induced (FeCl_3_) post-traumatic epilepsy (PTE) and lithium-pilocarpine-induced (LiCl-Pilo) status epilepticus (SE). By integrating Electrophysiology (EEG), γ-oscillation spectral analysis, multi-unit activity (MUA) recordings with behavioural assessments Morris Water Maze (MWM) and Open Field Test (OFT) we validated and established the chronic hyperexcitability and associated cognitive-emotional co-morbidities of these two models. Which further served as a functional backdrop for investigating a targeted 8-gene panel consisting *of GABRA2, KCNN2, KCNAB1, GRIK1, BCL11A, BRD7, STX1B,* and *PNPO*. RT-PCR analysis demonstrated a systemic dysregulation of 5 genes out of these 8 genes, mirroring the simultaneous collapse of inhibitory signalling, membrane stability, and neurotransmitter metabolism seen in human phenotypes. While the clinical roles of these genes are well documented, our study demonstrates that these rat models provide a high-fidelity platform for complex multigene studies and gene-level therapeutic exploration. Furthermore, in-silico docking identified Fisetin as a high-affinity ligand for the dysregulated proteins, offering a promising multi-target strategy for stabilizing the epileptic genome.

## Introduction

Epilepsy is a complex neurological dysfunction characterized by an enduring predisposition to generate spontaneous seizures, often accompanied by debilitating cognitive, psychological, and social comorbidities[1]. Affecting over 50 million people globally, it remains one of the most prevalent neurological disorders, with an estimated incidence of 4–10 per 1,000 individuals [2]. Despite the expansion of the antiepileptic drug (AED) arsenal and advances in surgery and neuromodulation, approximately one-third of patients remain refractory to treatment, suffering from uncontrolled seizures that significantly increase the risk of psychosocial stigma and mortality [3]. The process of epileptogenesis involves a transition where normal neural circuits are transformed into hyperexcitable networks, primarily through a fundamental imbalance between excitatory and inhibitory neurotransmission [4]. Acquired epilepsy, resulting from insults such as traumatic brain injury (TBI), stroke, or infections, represents a significant portion of clinical cases [5]. Post-traumatic epilepsy (PTE) is a frequent and lifelong complication of traumatic brain injury (TBI), often escalating into Status Epilepticus (SE) an extreme form of prolonged seizure activity with a mortality rate of approximately 20% [6]. The mechanism driving PTE involves the haemolysis of extravasated blood, leading to iron deposition within the brain parenchyma. This release of ferrous and ferric ions (Fe^2+^/Fe^3+^) triggers the Fenton and Haber–Weiss reactions, generating reactive oxygen species (ROS) that induce chronic oxidative stress, inflammatory responses, and ionic pathway dysfunction [7,8]. In the present study, we utilized the FeCl_3_-induced rat model to mimic human PTE, alongside the LiCl-Pilo model to simulate the systemic cholinergic overstimulation characteristic of SE [9]While the spectrum of human epilepsies is increasingly linked to genetic mutations covering over 23 distinct phenotypes most experimental research remains confined to single-gene models [10]. However, common forms of epilepsy are inherently multifactorial, involving a complex genetic architecture that transcends single-gene discovery[11,12]. There is a critical need to investigate these conditions at a multigene level to understand how various regulatory systems collapse simultaneously. Based on genome-wide association studies, we prioritized 8 genes, these include four ion channel genes (*GABRA2, KCNN2, KCNAB1, and GRIK1*) one transcription factors (*BCL11A*), the histone modification gene *(BRD7*), the synaptic transmission gene (*STX1B*) and the pyridoxine metabolism gene (*PNPO*) [13].Out of these only 5 genes showed systemic downregulation in the mRNA expression therefore were taken further in this study.

Furthermore, given the limitations of current AEDs, there is significant interest in natural bioactive compounds for their neuroprotective and antioxidant capabilities. Curcumin, derived from Curcuma longa, and quercetin, a ubiquitous flavonoid, have been extensively investigated for their ability to modulate ion channels and suppress neuroinflammation[14,15]. Alongside these, Fisetin has emerged as a potent antioxidant and neuroprotective agent [16]. While all three compounds show promise, it is imperative to investigate their impact at the gene-expression level within a multigene framework. In this study, we utilized protein-ligand docking to evaluate the binding affinities of curcumin, quercetin, and fisetin against the protein structures of 3 genes only as the alpha-fold structure was available for only these 3 genes *GABRA2, KCNN2, and PNPO*. Interestingly, while all compounds demonstrated interaction, fisetin consistently exhibited the most robust binding energy and stability, particularly across the GABRA2, KCNN2, and PNPO proteins. This superior docking performance suggests that Fisetin may act as a more effective multi-target genomic stabilizer than curcumin or quercetin. By bridging these in-silico predictions with in-vivo expression analysis of the five-gene panel, this study seeks to provide new insights into the pathophysiology of acquired epilepsy and establish these rat models as high-fidelity platforms for gene-level therapeutic exploration.

## Materials and Methods

### Materials

Pilocarpine hydrochloride, FeCl_3_, lithium chloride (LiCl), scopolamine methyl bromide, and TRI reagent were procured from Sigma Aldrich (USA). Diazepam and isoflurane were obtained from Lori (India) and Neon (India), respectively. For electrophysiological recordings, tissue-compatible stainless-steel screw electrodes and bipolar electrodes were sourced from Plastics One (USA). Molecular biology reagents, including the RevertAid™ First Strand cDNA Synthesis Kit and EvaGreen™ qPCR Mastermix, were purchased from Thermo Fisher Scientific (Lithium) and G-Biosciences (USA). All additional reagents were of analytical grade.

### Animals and Experimental Design

Male Wistar rats (250–300 g) were obtained from the Central Laboratory Animal Research (CLAR) at Jawaharlal Nehru University (JNU), New Delhi. Animals were housed in polypropylene cages under a controlled 12:12 h light-dark cycle and ambient temperature with ad libitum access to food and water. All experimental protocols were approved by the Institutional Animal Ethics Committee (IAEC Code: 14/2021) and performed in accordance with the Committee for the Purpose of Control and Supervision of Experimental Animals (CPCSEA) guidelines.

### A total of 40 rats were randomized into four groups (n=10 per group)

1. **Saline Control:** Received intracortical saline; served as the control for the PTE model.
2. **FeCl_3_ (PTE):** Received intracortical 100 mM FeCl_3_
3. **Saline Control**: Received intraperitoneal (i.p.) saline; served as the control for the SE model.
4. **LiCl-Pilo (SE):** Received LiCl (127 mg/kg, i.p.) followed by Pilocarpine (125 mg/kg, i.p.).

### Surgical Procedures and Model Induction

For the PTE model, rats were anesthetized with 4% isoflurane and placed in a stereotaxic frame. A 5 µl volume of 100 mM FeCl_3_ was microinjected into the somatosensory cortex (AP: -1.0 mm, ML: 1.0 mm, DV: -1.0 mm relative to bregma) over 5 min. For Status Epilepticus (SE) induction, rats were pre-treated with LiCl (127 mg/kg, i.p.) 23 h prior to induction. Scopolamine methyl bromide (1 mg/kg, i.p.) was administered 30 min before Pilocarpine (125 mg/kg, i.p.) to mitigate peripheral cholinergic effects. Seizure severity was monitored using a modified Racine Scale; only rats exhibiting Stage 4 (rearing), 5 (rearing and falling), or 6 (tonic-clonic seizures) were included. To reduce mortality, Diazepam (8 mg/kg, i.p.) was administered 120 min post-SE onset.

#### Electrode Implantation

In all groups, epidural screw electrodes were implanted (AP: 2.0 mm, ML: 2.0 mm), with a bipolar electrode targeted at the hippocampus (AP: -2.8 mm, ML: 2.5 mm, DV: -2.71 mm). A reference electrode was placed at the frontal sinus. Electrodes were secured with dental acrylic and soldered to a nine-pin connector. Rats were allowed 8 days for recovery.

### EEG and MUA Recordings

Recordings were conducted on the 30^th^ day post-induction in awake, immobile rats after a 4- day habituation period. Extracellular signals were amplified (P511, Grass Technologies) and filtered (EEG: 1–100 Hz; MUA: 300 Hz–10 kHz). MUA counts were isolated using a window discriminator (WP1 Model 121) and quantified alongside EEG traces using PolyVIEW16 software. *Spectral Power Analysis*: Synchronous gamma oscillations (30–50 Hz) were analysed using Fast Fourier Transform (FFT) with a Hanning window (10s length). Power spectral density (Vrms^2/Hz) was calculated, and relative power was expressed as a percentage of total power to minimize inter-subject variability.

### Behavioral Assessments

#### Morris Water Maze (MWM)

Spatial learning was assessed from days 26–30^th^. Rats were trained to find a submerged platform (diameter: 15 cm) in a circular pool (diameter: 168 cm). Escape latency (time to reach the platform) was recorded over four trials per day (max 60 s).

#### Open Field Test (OFT)

On day 30^th^, exploratory behaviour and anxiety were measured in a 16-square arena. Parameters included ambulatory activity (squares crossed), rearing (vertical movements), and the defecation index over a 5 min period.

### Gene Expression Analysis

Cortical and hippocampal tissues (n=5) were harvested on day 30^th^. Total RNA was extracted using TRI-Reagent and quantified (NanoDrop 2000). 1 µl of RNA was reverse-transcribed (RevertAid Kit). Quantitative Real-Time PCR (qRT-PCR) was performed for the 5-gene panel (*GABRA2, KCNN2, KCNAB1, STX1B,* and *PNPO)* using EvaGreen™ Mastermix. GAPDH served as the internal control. Fold changes were determined using the 2T‒ΔΔC method. Fold change = 2T-ΔΔCT, where ΔΔCT = (CT, target gene-CT, GAPDH) control – (CT, target gene- CT, GAPDH) experimental.

### In-Silico Molecular Docking

#### Ligand/Protein Preparation

3D structures for Curcumin, Fisetin, and Quercetin were retrieved from PubChem and prepared via Schrödinger’s LigPrep (OPLS_2005 force field). Target protein structures for GABRA2 (PDB: 6DW0), KCNN2 (PDB: 3SJQ), and PNPO (AlphaFold) were retrieved and validated using Ramachandran plots (PROCHECK)[17,18].

#### Docking & Binding Energy

Proteins were optimized using the Protein Preparation Wizard. Receptor grids were generated around identified binding pockets (e.g., ABU site for GABRA2; Ca^2+^ site for KCNN2). Molecular docking was performed using the GLIDE module (HTVS, SP, and XP modes). The binding free energy was calculated using the Prime MM-GBSA module to determine the stability of the protein-ligand complexes[19].

### Statistical Analysis

Data are presented as Mean ± SEM. Significance was determined using Student’s t-test for MUA, spectral power, and RT-PCR. MWM escape latency was analysed via two-way ANOVA with Sidak’s post-hoc test. A p-value < 0.05 was considered statistically significant.

## Results

### Electrophysiological Validation

#### EEG Paroxysms and MUA count on day 30^th^ day of post-induction

chronic epileptiform activity was confirmed in both the cortical and hippocampal regions of FeCl_3_ and LiCl-Pilo- injected rats. EEG recordings revealed hallmark seizure signatures, including spike-wave complexes, solitary spikes, and sharp wave paroxysms (Fig. 1A-B). To ensure these signals reflected local cellular hyperexcitability, we recorded concurrent MUA. In the FeCl_3_ model, MUA counts increased significantly in the cortex (p≤0.001) and hippocampus (p≤0.01) compared to controls (Fig. 1C). Similarly, the LiCl-Pilo model exhibited a robust increase in MUA counts (p≤0.001 and p≤0.001) across both regions (Fig. 1D), providing functional evidence of an established epileptic network.

**Fig 1:**
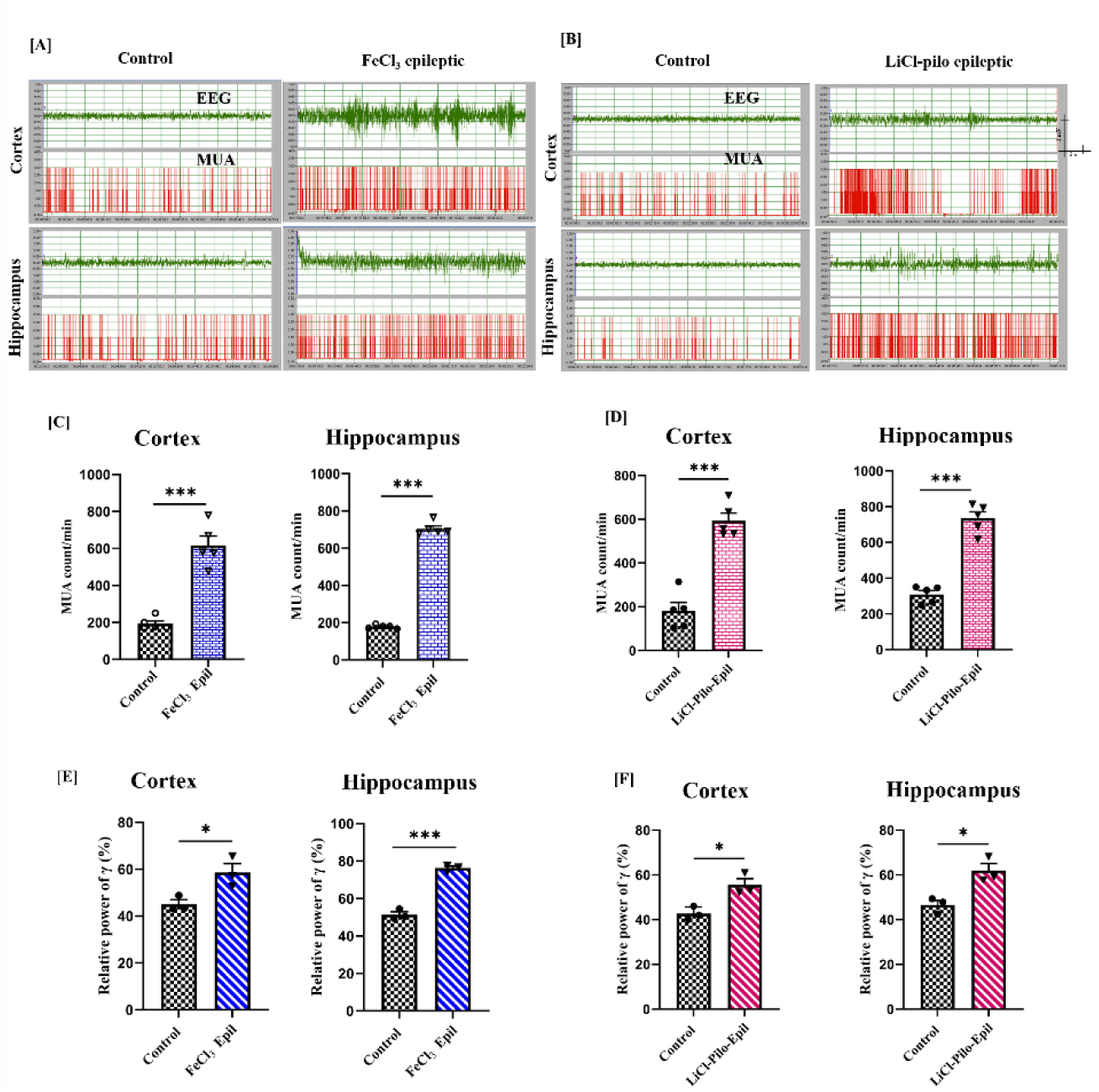
EEG, MUA counts and spectral power analysis of γ-oscillations in the cortex and hippocampus of experimental rats. Representative EEG stretches of 1 min duration obtained after 30th day of stereotaxic surgery in FeCl3 (A) and LiCl-Pilo (B) epileptic rats. Bar graphs showing MUA counts in the cortex and hippocampus of FeCl3 (C) and LiCl-Pilo (D) epileptic rats. Relative spectral power (%) of γ-oscillation from the cortex and hippocampus of FeCl3 (E) and LiCl-Pilo (F) epileptic rats. The data for the MUA counts are expressed as the means ± SEM (n = 5), and the relative mean spectral power is expressed as the mean ± SEM (n = 3). *p ≤ 0.05, **p ≤ 0.01 and ***p ≤ 0.001, significantly different from the control group.

**Table 1.** **A.** The docking results for the AlphaFold GARBA2 protein with ligands (curcumin, quercetin, and fisetin) representing compound name, PubChem ID, docking scores, binding energies and interacting residues with their rank.

| Rank | Name | PubChem ID | Docking score (kcal/mol) | Binding energy (kcal/mol) | Interacting residues with 5 Å |
| --- | --- | --- | --- | --- | --- |
| 1 | Fisetin | 5281614 | -4.891 | -41.01 | N130, Y237, Y187, V239, T234, S232, S186, I230, H129, and F127. |
| 2 | Quercetin | 5280343 | -4.861 | -42.36 | S186, Y237, Y187, V239, T234, S232, I230, G185, F127, H129, and A188. |
| 3 | Curcumin | 969516 | -3.277 | -34.1 | T190, Y237, Y187, V239, T234, S233, S232, S186, I230, H129, G185, F127, A188. |

| Rank | Name | PubChem ID | Docking score (kcal/mol) | Binding energy (kcal/mol) | Interacting residues with 5 Å |
| --- | --- | --- | --- | --- | --- |
| 1 | Fisetin | 5281614 | -3.85 | -26.52 | N53, V55, R74, E54, D56, A57, and K75 from chain A; A480 from chain C. |
| 2 | Quercetin | 5280343 | -3.715 | -25.45 | N53, V55, R74, E54, D56, A57, K75 and D78 from chain A; N477 and A480 from chain C. |
| 3 | Curcumin | 969516 | -3.24 | -30.23 | N53, V55, T70, R74, K75, E67, E54, D56, A57, and M71 from chain A; L476, N477, and A480 from Chain C. |

| Rank | Name | PubChem ID | Docking score (kcal/mol) | Binding energy (kcal/mol) | Interacting residues with 5 Å |
| --- | --- | --- | --- | --- | --- |
| 1. | Fisetin | 5281614 | -5.465 | -52.77 | N112, Q174, Y157, W206, T111, R161, L98, K117, K100, and E77. |
| 2. | Quercetin | 5280343 | -5.55 | -48.94 | N112, Y157, W206, T111, R161, Q174, L98, K100, E77, F110, and A169. |
| 3 | Curcumin | 969516 | -4.992 | -53.68 | N112, Y205, Y157, W206, V170, T111, S172, S165, R173, R161, R161, Q174, P203, P162, L98, K100, G168, F110, and A169. |
| 4. | PLP | 1051 | -7.6 | -20.71 | Y157, A169, F110, G168, L97, L98, N112, P162, Q174, R161, S165, T111, W206, W206, Y157, and K100. |

### Spectral Power Analysis

Pathological gamma oscillations spectral power analysis was utilized to quantify synchronous network oscillations. In FeCl_3_-treated rats, relative gamma power (30–50 Hz) was significantly elevated in the cortex (p≤0.05) and hippocampus and (p≤ 0.001) (Fig. 1E). This pathological shift was mirrored in the LiCl-Pilo model (p≤0.05) for both regions), indicating that despite different primary triggers, both models converge on a shared frequency-domain signature of hyperexcitability (Fig. 1F).

### Behavioral Comorbidities

Cognitive impairment and anxiety spatial learning: The MWM test revealed profound spatial memory deficits. FeCl_3_-induced rats showed significantly prolonged escape latencies from day 3 through day 5 (p ≤0.05) (Fig. 2A). In the LiCl-Pilo group, these deficits appeared earlier (day 2) and reached a high significance by day 5 (p ≤ 0.01) (Fig. 3A). In the OFT, exploratory behaviour in both models demonstrated a significant reduction in ambulatory and rearing activities (p ≤ 0.001), coupled with a sharp increase in the fecal boli index (p ≤0.01) (Fig. 2B- D, 3B-D). Together, these metrics confirm a state of high anxiety and reduced motor motivation associated with chronic epilepsy.

**Fig 2:**
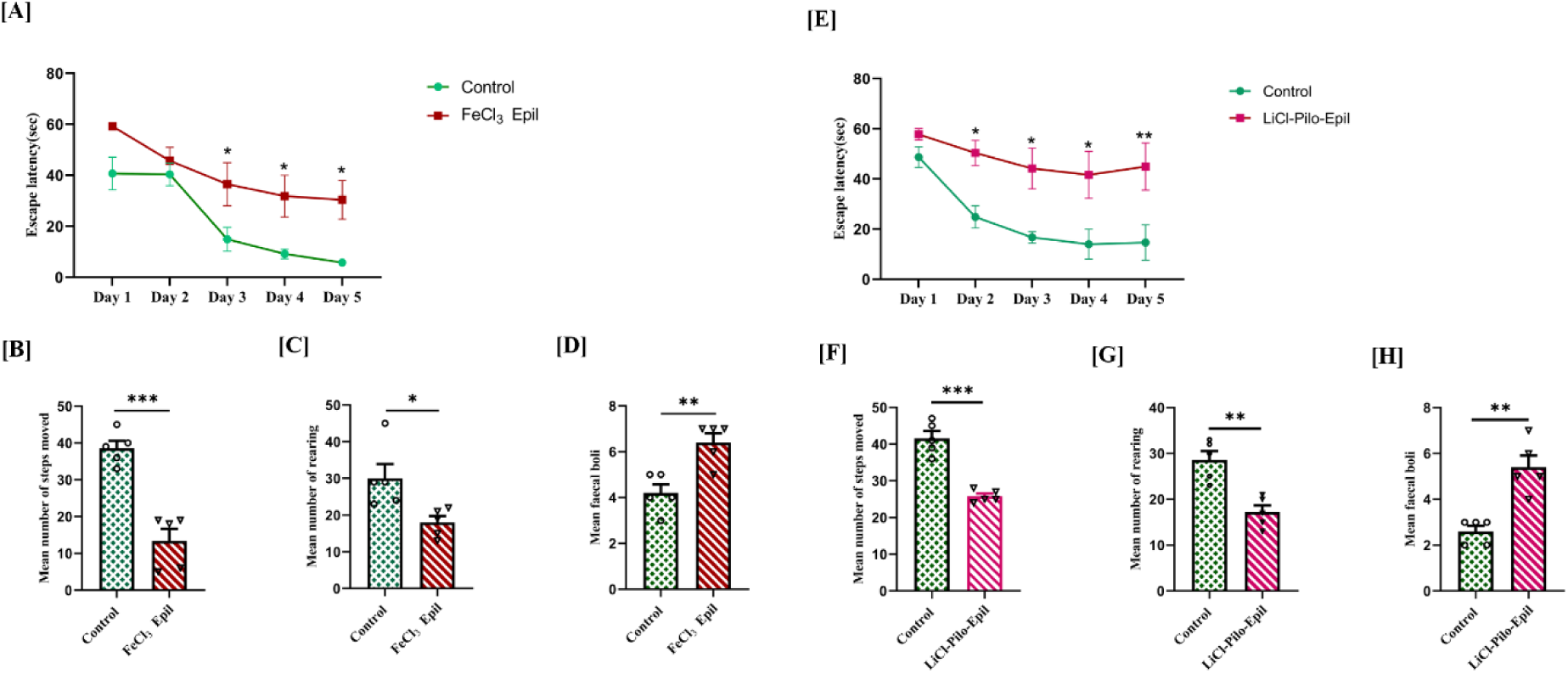
Behavioral analysis of control and FeCl3-induced epileptic rats. Escape latency to reach the hidden platform (A&E), ambulatory activity (B&F), rearing activity (C&G) and the fecal index (D&H) in control and epileptic rats. The data are expressed as the means ± SEMs (n=5). *p ≤ 0.05, **p ≤ 0.01 and ***p ≤ 0.001 indicate significant differences from control rats.

**Fig 3:**
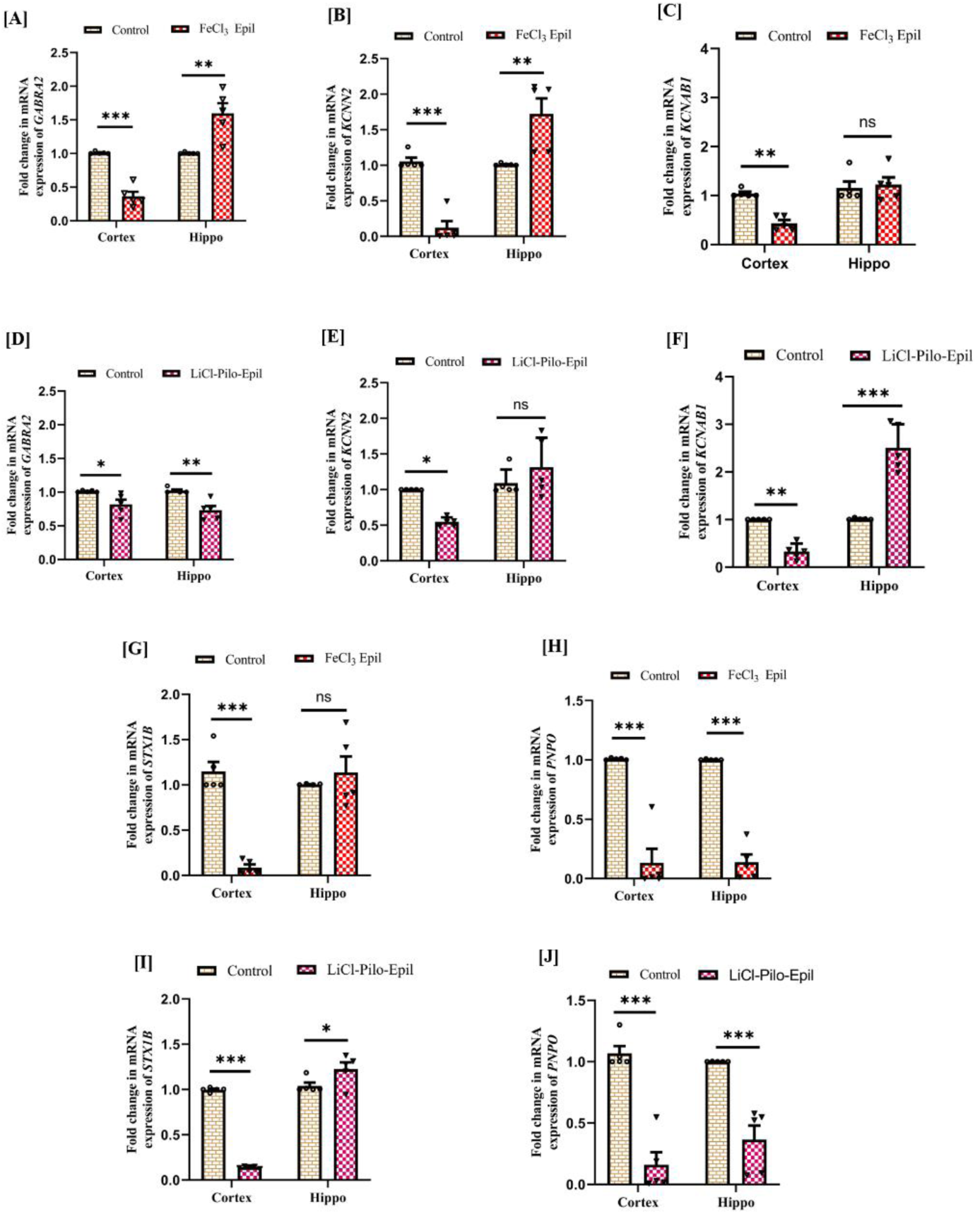
mRNA expression of epilepsy-associated ion channel genes in the cortex and hippocampus of experimental rats. The bar graphs show the fold changes in the mRNA expression of *GABRA2* (A, D), *KCNN2* (B, E), *KCNAB1* (C, E), *STX1B (*G, I) and *PNPO* (H, J) in the cortex and hippocampus of FeCl3 and LiCl-Pilo-induced epileptic rats. The data are expressed as the means ± SEM (n=5). *p ≤ 0.05, **p ≤ 0.01 and ***p ≤ 0.001. This value significantly differed from that of the control rats.

### Multigene Expression Profiles

The mRNA expression of the prioritized 5-gene panel revealed a complex regulatory collapse. In the cortex, both models showed a significant downregulation of *GABRA2, KCNN2*, and *KCNAB1* (p≤0.05 to p≤0.001) (Fig.3 A-F) explaining the loss of inhibitory control and membrane stability. Interestingly, we observed a compensatory or maladaptive upregulation of these genes in the hippocampus of FeCl_3_ rats, while LiCl-Pilo rats showed a significant decrease in *GABRA2* (p ≤0.001) but an increase in *KCNAB1* (Fig. 3). Furthermore, *STX1B* (synaptic transmission) was consistently downregulated in the cortex of both models (p ≤0.001) (Fig. G&I), while *PNPO* (metabolism) was severely suppressed across all regions and both models (p ≤0.001), indicating a systemic failure in neurotransmitter synthesis (Fig. 3 H&J).

### In-Silico Validation

Fisetin as a multi-Target lead protein structure validation via PROCHECK confirmed that 86% of residues in our target proteins (GABRA2, KCNN2, and PNPO) resided in favorable regions (Fig. S1 & S2). **GABRA2 Docking**: Fisetin emerged as the top-ranked ligand with the best docking score (-4.891 kcal/mol) and strong binding energy (-41.01 kcal/mol) (Fig. 4A), outperforming curcumin (Fig. 4B) and quercetin (Fig. 4C) in orientation stability (Table. 1A). **KCNN2 & PNPO Docking**: Fisetin again showed the most favourable docking score for KCNN2 (-3.85 kcal/mol). While curcumin showed high binding energy for PNPO, fisetin maintained a highly significant binding energy (-52.77 kcal/mol) and the most favourable pose for multi-target stabilization (Fig. 4D & G) (Table. 1 B & C).

**Fig 4:**
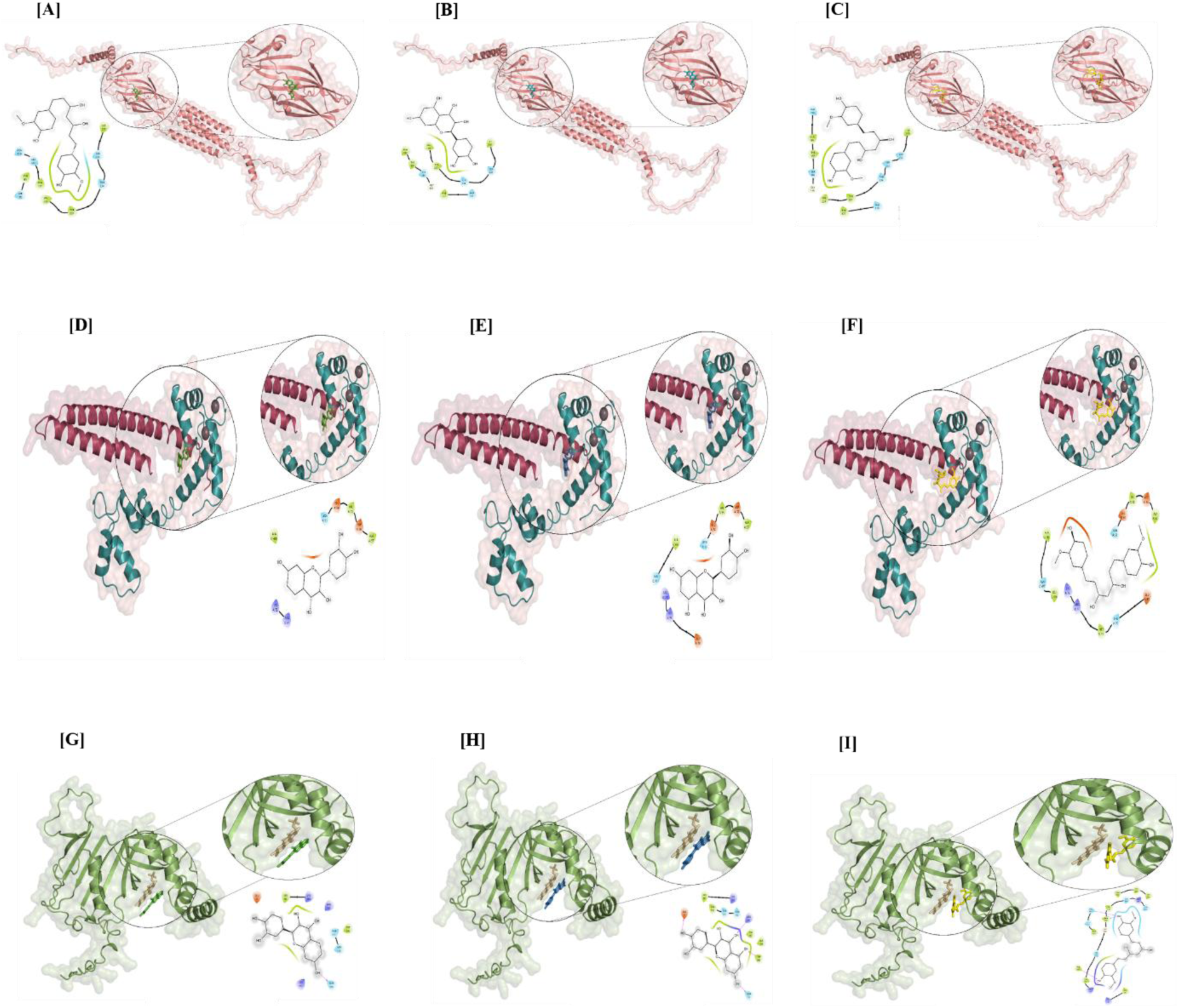
3D structure of docked complex AF-GABR2 protein chain A colored in salmon interacting with fisetin_5281614 as a green stick (center) (A), quercetin_5280343 as a blue stick (center) (B) and curcumin_969516 as a yellow (center) (C) with the GABRA2 protein in its active site (bottom left). The binding poses of fisetin in green, quercetin in sky blue and curcumin in yellow. fisetin_5281614 in a green stick (left) (D), quercetin_5280343 in a blue stick (left) (E) and curcumin_ 969516 in a yellow (left) (F) interact with protein KCNN2 in the active site (bottom right). The docked complex AF-PNPO FMN bound protein colored in smudge-green in cartoon representation and FMN colored in purple sticks interacting with) fisetin_5281614 in green sticks (left) (A), quercetin_5280343 in sky blue sticks (left) (B) and curcumin_969516 in yellow sticks (left) (C).

## Discussion

The results of the present investigation provide a comprehensive comparative analysis of two distinct models of epilepsy, the FeCl_3_-induced post-traumatic epilepsy (PTE) model and the lithium-pilocarpine (LiCl-Pilo) induced status epilepticus (SE) model uncovering significant parallels in electrophysiological disruption, Behavioral morbidity, and genomic instability. By evaluating Racine scores alongside cortical and hippocampal EEG recordings, our study validates the induction of chronic hyperexcitability in both experimental groups. In the FeCl_3_ model, the focal injection of iron into the cerebral cortex induces a persistent epileptogenic focus through oxidative stress-mediated mechanisms. As initially described by Willmore et al. (1978), this model serves as a robust surrogate for human post-traumatic epilepsy, where reactive oxygen species (ROS) attack neural membranes, leading to lipid peroxidation[20]. Our findings suggest that this oxidative environment disrupts the astrocytal absorption of glutamic acid, likely due to the reduction of glial glutamate transporter proteins [21]. The resulting accumulation of extracellular glutamate facilitates epileptogenesis, creating a network of hyperexcitability that originates in the cortex and spreads to subcortical structures like the hippocampus. This transmission is likely mediated by the entorhinal-perirhinal-cortical connections, a finding that aligns with previous research on hippocampal-cortical networks in seizure propagation [22]. The synchronization of these networks is fundamental to the transition from a focal insult to a generalized seizure state, a phenomenon we observed clearly in the spreading of epileptiform discharges in our cortical recordings.

Conversely, the LiCl-Pilo model represents a different pathological trajectory characterized by intense cholinergic stimulation. The administration of lithium chloride sensitizes muscarinic receptors and increases acetylcholine levels in the hippocampus [23]. The subsequent pilocarpine-induced SE triggers widespread neuronal damage, particularly in the piriform cortex, amygdala, and the hilus of the dentate gyrus [24]. Our observation of spontaneous recurrent seizures (SRS) following a latent phase confirms the formation of new, hyperexcitable circuits that persist throughout the life of the animal[25]. The EEG patterns in these rats showed both interictal spikes and ictal polyspikes, mirroring the clinical progression seen in human acquired epilepsy [26]. A critical convergence between both models was the significant increase in gamma (γ) oscillation power (30-50 Hz) revealed through spectral analysis. Gamma rhythms are essential for higher-order cognitive functions and inter-regional communication, but their pathological elevation often precedes seizure onset[27]. Our results support the theory that sudden, abnormal gamma activity reflects a malfunction in the inhibitory interneuron network, often brought on by oxidative stress and neuroinflammation[28]. These fast rhythms likely reflect the neural network’s production of ictal-like discharges, a process that gradually increases in frequency from the pre-ictal to the ictal state. The Behavioral co-morbidities observed in our study, specifically the cognitive deficits in the MWM and the anxiety-like behaviour in the OFT, further underscore the severity of the induced conditions. The increased latency to reach the hidden platform in MWM-tested rats indicates a failure in hippocampal-dependent spatial learning, which is a common clinical complaint in patients with temporal lobe or post-traumatic epilepsy [29,30]. Furthermore, the reduction in exploratory rearing and the increased fecal index in the OFT suggest a significant decline in motor motivation and an increase in emotional distress. These Behavioral markers correlate with the extent of cellular impairment observed in the CA1 region of the hippocampus, which our MUA recordings confirmed as a primary site of epileptiform activity [31]. By linking these behavioral outcomes to the electrophysiological data, we establish that the “epileptic brain” in these rats is not only prone to seizures but is fundamentally altered in its ability to process information and regulate emotion. The inclusion of MUA counts in our study proved essential, as it ensured that the recorded signals were true reflections of cellular activity rather than unintentional volume conductance from adjacent structures [32].

A primary objective of this research was to correlate these physiological and behavioral changes with the expression of key genes associated with human epilepsy. While the individual roles of these genes are well-established in the context of human clinical epilepsy, their systemic interplay during the progression of the disease remains poorly understood. Our study provides significant insights by demonstrating that these rat models can be utilized as high-fidelity platforms for complex multigene studies. By capturing the simultaneous dysregulation of these pathways, this work offers a novel framework for exploring targeted gene-level interventions and evaluating therapeutic efficacy in a way that clinical observation alone cannot achieve. This is particularly relevant for exploring the “genomic landscape” of the disease and helping in the development of treatment methods at the gene level. Our analysis focused on five specific candidate genes, discussed hereafter in the order of their functional contribution to the hyperexcitable state: *GABRA2, KCNN2, KCNAB1, STX1B, and PNPO*.

Regarding *GABRA2*, which encodes the GABA_A receptor α2 subunit protein, we observed a significant dysregulation that mirrors human epileptic syndromes. As the primary inhibitory neurotransmitter system in the brain, GABA maintains the vital balance between excitation and inhibition. Defects in *GABRA2* expression are known to lower the amplitude of GABA-evoked potentials, leading to various epileptic syndromes and intellectual disabilities [33]. Our results showed a general downregulation of *GABRA2* mRNA in the cortex and hippocampus of both models, which likely contributes to the loss of inhibitory control. Interestingly, an upregulation was noted specifically in the hippocampus of the FeCl_3_-induced rats. This localized increase may represent an unsuccessful or maladaptive compensatory response to the focal iron-induced excitation. Such inconsistencies in *GABRA2* expression have been noted in other neurological conditions like schizophrenia, suggesting that the precise regulation of this gene is essential for maintaining neural network oscillations [34]. The disruption of *GABRA2* directly correlates with our spectral power analysis, as altered GABAergic signalling is a known driver of abnormal gamma oscillations [35].

Following the disruption of inhibitory signalling, we investigated the potassium channel-related genes *KCNN2* and *KCNAB1*, both of which are critical for maintaining membrane stability and repolarization. The *KCNN2* gene encodes the SK2 subunit of calcium-activated potassium channels, which govern the firing patterns and excitability of central neurons. We found a significant decline in *KCNN2* expression across the epileptic models, a result that aligns with the observed behavioral tremors and locomotor instability in our rats [36]. This downregulation reduces the after-hyperpolarization (AHP) effect, leading to the neuronal hyperexcitability and hereditary seizure vulnerability seen in our EEG. Parallel to this, the *KCNAB1* gene, which encodes a voltage-gated potassium channel beta-1 subunit, was significantly downregulated in the cortex of both models. Given that *KCNAB1* deletions in mice are linked to impairments in associative learning and amygdala hyperexcitability, this genetic decline provides a molecular basis for the cognitive impairments observed in our behavioral studies [37,38]

The investigation into synaptic transmission involved the *STX1B* gene, which encodes syntaxin-1B, a protein essential for synaptic vesicle release. Abnormal synaptic transmission is a significant driver of seizure onset and progression. In our study, mRNA downregulation of *STX1B* was reported in the cortex of both the FeCl_3_ and LiCl-Pilo models. However, an increase in mRNA expression was noted in the hippocampus of LiCl-Pilo rats. This divergence suggests that different brain regions may respond differently to the same systemic insult, with some areas undergoing a loss of synaptic integrity while others may experience an aberrant increase in transmitter release [39,40] Such mis regulation of the synaptic machinery facilitates the chaotic firing patterns observed in our EEG analysis.

Finally, we evaluated the *PNPO* gene, which is essential for vitamin B6 metabolism and the synthesis of neurotransmitters such as GABA, dopamine, and serotonin. We found a significant decline in *PNPO* expression in the cortex and hippocampus across both models. This metabolic failure likely leads to a shortage of the PLP cofactor, resulting in resistant seizure patterns and neonatal-like epileptic encephalopathy phenotypes observed in clinical settings[41,42]. Because *PNPO* is responsible for the production of several neurotransmitters that decrease neuronal excitability, its downregulation represents a “bottleneck” in the brain’s ability to defend itself against epileptogenesis [43]. By connecting these five genes, we demonstrate a multi-level collapse of the brain’s regulatory systems, spanning from receptor inhibition *(GABRA2)* and membrane stability *(KCNN2, KCNAB1*) to synaptic release *(STX1B)* and neurotransmitter metabolism *(PNPO)*.

Furthermore, our in-silico docking analysis successfully identified Fisetin as a superior multi-target lead compound, exhibiting the most stable binding affinities across our prioritized gene panel compared to curcumin and quercetin. Ultimately, this work establishes that acquired epilepsy is not merely a disorder of abnormal firing, but a systemic genomic failure. By bridging functional electrophysiology with genomic instability and computational drug discovery, we offer a novel platform for the development of multi-target therapies for preventing the entrenchment of epilepsy and mitigating its long-term neurological consequences.

## Conclusion

In conclusion, our study demonstrates that while the initiating events of the FeCl_3_ and LiCl- Pilo models differ one being localized trauma and the other systemic cholinergic overstimulation they culminate in a shared pathological state. This state is defined by abnormal gamma oscillations, profound cognitive and emotional dysfunction, and a specific molecular signature involving the dysregulation of *GABRA2, KCNN2, KCNAB1, STX1B*, and *PNPO*. Our in-silico docking studies further suggest that natural bioactive compounds like Fisetin, which showed high binding affinity for GABRA2, KCNN2, and PNPO, may offer a promising multi-target therapeutic strategy. Unlike traditional mono-therapies, these compounds may stabilize the brain’s genetic and metabolic landscape, providing a more holistic approach to treating the complex reality of human epilepsy. This study provides a comprehensive comparative analysis of two distinct rat models of epilepsy, demonstrating a convergent pathway of genomic instability and molecular dysregulation. By utilizing a high-fidelity multigene framework, we have shown that both post-traumatic (FeCl_3_) and chemical (LiCl-Pilo) insults culminate in a shared pathological phenotype characterized by chronic gamma hyperexcitability, cognitive-emotional comorbidities, and the systemic collapse of key ion channel and metabolic pathways specifically involving *GABRA2, KCNN2, KCNAB1, STX1B, and PNPO*.

## Ethics declarations

All experimental protocols carried out in rats were approved by the Institutional Animal Ethics Committee (IAEC Code: 14/2021) were obtained from the Central Laboratory Animal Research (CLAR) at Jawaharlal Nehru University (JNU), New Delhi and performed in accordance with the Committee for the Purpose of Control and Supervision of Experimental Animals (CPCSEA) guidelines.

## Acknowledgments

The authors acknowledge the Central Instrument Facility, School of Life Sciences and Central Laboratory Animal Resources, Jawaharlal Nehru University, New Delhi, for facilitating the experiments.

## Data availability

The data will be available upon a reasonable request.

