## Supplementary material for "Expression Pattern of Epilepsy Associated Genes as a Common Pathological Signature in PTE and SE models and possible therapeutics via an in-silico approach": Supplimentary table S1

**Table S1.** List of primer sequences used for the RT-PCR analysis.

| Genes | Primer sequences | Product length (bp) |
|---|---|---|
| *GABRA2* | Forward 5'- CCAGTCAATTGGGAAGGAGA-3'<br>Reverse 5'-CGCGTAACAAACAGCGATAA-3' | 160 |
| *KCNN2* | Forward 5'- TCTTGATTGCCAGAGTCATG-3'<br>Reverse 5'- GCTCAGAAGTCGGTCCTCAC-3' | 161 |
| *KCNAB1* | Forward 5'- GAAGATTGGGGACGCTGATA-3'<br>Reverse 5'- AGTTGAAAGCAAGCCCAAGA-3' | 175 |
| *STX1B* | Forward 5'- CGACACCAAGAAAGCTGTGA-3'<br>Reverse 5'- TGCTATCTATGTGCGGCAAG-3' | 169 |
| *PNPO* | Forward 5'- GGCTCATCTGACCTCTCTCTGG-3'<br>Reverse 5'- TCTCCGGCAGTTTCTTCACT-3' | 198 |
| *FOXO3a* | Forward 5'- GACCTGCTCACTTCGGACT-3'<br>Reverse 5'- TTTGCATAGACTGGCTGACG-3' | 163 |
| LINE-1 ORF1 | Forward 5'- AGCGAGGATGTGGAGAAAGA-3'<br>Reverse 5'- GCTGCGATGAACATAGTGGA-3' | 192 |
| LINE-1 ORF2 | Forward 5'- GCTCCTCATGGCACTTTCTC-3'<br>Reverse 5'- TGGGCATTCTTCCCTTATTG-3' | 198 |
| *GAPDH* | Forward 5'- AACGACCCCTTCATTGAC-3'<br>Reverse 5'- TCCACGACATACTCAGCAC-3' | 191 |

**Fig. S1:**

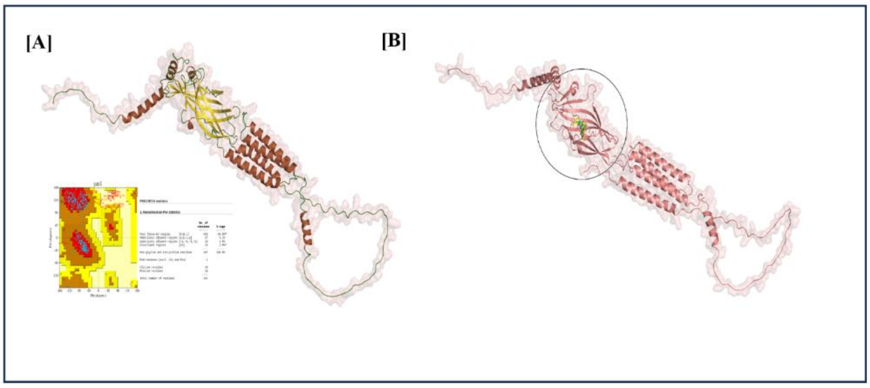


3D structure of alphaFold GABRA2 protein (A) showing a helix, beta sheets, and loop in smudge-green, cyan, and red, along with the PROCHECK Ramachandran plot for the validation of the AlphaFold-GABRA2 protein. 3D structure of docked complex AF-GABAR2 ligands; the GABAR2 protein chain A is colored in salmon in cartoon representation, and all three ligands are represented as sticks, where fisetin is in green, quercetin is in teal, and curcumin is in yellow (B).

**Fig. S2:**

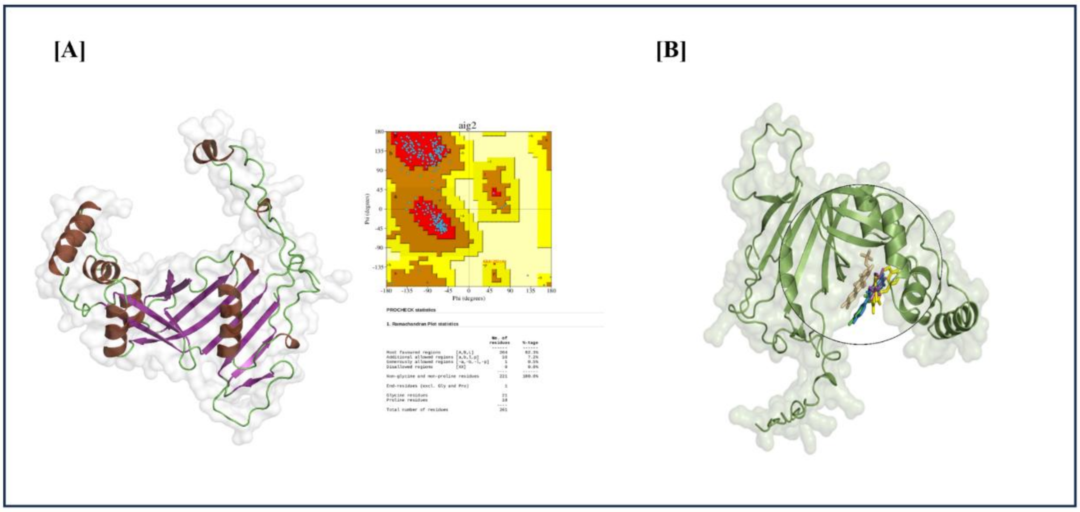


3D structure of alphaFold PNPO protein, where the helix, beta sheets, and loop are colored brown, slate blue, and green (left), along with the PROCHECK Ramachandran plot for the validation of the AlphaFold-PNPO protein with statistics (right) (A). Superimposition of the ligands (all three compounds) in the binding pocket of the docked complex AF-PNPO-Ligands; PNPO protein chain A is colored smudge-green, FMN is colored purple, and the ligands (all poses) are colored stick. The binding modes of PLP are shown in magenta, quercetin in sky blue, fisetin in green, and curcumin in yellow (B).
